# S1PR3 mediates itch and pain via distinct TRP channel-dependent pathways

**DOI:** 10.1101/235614

**Authors:** Rose Z. Hill, Takeshi Morita, Rachel B. Brem, Diana M. Bautista

## Abstract

Sphingosine 1-phosphate (S1P) is a bioactive signaling lipid associated with a variety of chronic pain and itch disorders. S1P signaling has been linked to cutaneous pain, but its role in itch has not yet been studied. Here we find that S1P triggers itch and pain in mice in a concentration-dependent manner, with low levels triggering acute itch alone, and high levels triggering both pain and itch. Calcium imaging and electrophysiological experiments revealed that S1P signals via S1PR3 and TRPA1 in a subset of pruriceptors, and via S1PR3 and TRPV1 in a subset of heat nociceptors. And in behavioral assays, S1P-evoked itch was selectively lost in mice lacking TRPA1, whereas S1P-evoked acute pain and heat hypersensitivity were selectively lost in mice lacking TRPV1. We conclude that S1P acts via different cellular and molecular mechanisms to trigger itch and pain. Our discovery elucidates the diverse roles that S1P signaling plays in somatosensation and provides insight into how itch and pain are discriminated in the periphery.

**Significance Statement:** Itch and pain are major health problems with few effective treatments. Here, we show that the pro-inflammatory lipid S1P and its receptor S1PR3 trigger itch and pain behaviors via distinct molecular and cellular mechanisms. Our results provide a detailed understanding of the roles that S1P and S1PR3 play in somatosensation, highlighting their potential as targets for analgesics and antipruritics, and provide new insight into the mechanistic underpinnings of itch versus pain discrimination in the periphery.

## Introduction

Sphingosine 1-phosphate (S1P) is a bioactive sphingolipid associated with a variety skin disorders, including psoriasis (Checa et al., 2015; Myśliwiec et al., 2016), atopic and allergic contact dermatitis (Kohno et al., 2004; Sugita et al., 2010), and scleroderma (Castelino and Varga, 2014), as well as neuropathic pain (Patti et al., 2008; Janes et al., 2014), and other inflammatory diseases (Rivera et al., 2008; Chiba et al., 2010; Trankner et al., 2014; Donoviel et al., 2015; Roviezzo et al., 2015). S1P signaling via S1P Receptor 1 (S1PR1) facilitates the migration, recruitment, and activation of immune cells (Matloubian et al., 2004; Cyster and Schwab, 2012), and S1P is also thought to influence epithelial cell differentiation and cell division via the action of several other S1PRs (Schüppel et al., 2008; Japtok et al., 2014). Intriguingly, expression of both S1PR1 and S1PR3 has been reported in the somatosensory ganglia (Camprubi-Robles et al., 2013; Usoskin et al., 2015), and S1P modulates sensory neuronal excitability (Zhang et al., 2006a, 2006b; Mair et al., 2011; Camprubi-Robles et al., 2013; Li et al., 2015; Wang et al., 2017), and has been implicated in pain and pain hypersensitivity (Mair et al., 2011; Camprubi-Robles et al., 2013; Finley et al., 2013; Hou and Fang, 2015; Weth et al., 2015; Hill et al., 2018). We and others have recently shown an important role for S1P signaling via S1P Receptor 3 in cutaneous pain sensation in rodents (Camprubi-Robles et al., 2013; Hill et al., 2018); however, the downstream mechanisms by which S1PR3 triggers neuronal excitation and pain are unclear. Likewise, it is not known whether S1P also acts as a pruritogen to cause itch.

S1P has been examined in a variety of inflammatory skin diseases and disease models. There is evidence supporting both protective and pathogenic effects of S1P signaling in mouse and human chronic itch and inflammatory diseases. In humans, elevated serum S1P is correlated with disease severity in both psoriasis and systemic sclerosis (Castelino and Varga, 2014; Checa et al., 2015; Thieme et al., 2017). Furthermore, ponesimod, an S1PR modulator and blocker of S1P signaling, appears to be promising for psoriasis treatment in humans (Brossard et al., 2013; Vaclavkova et al., 2014; Krause et al., 2017). In contrast, in the imiquimod mouse model of psoriasis, it was observed that topical S1P exerts protective effects (Schaper et al., 2013), and in dogs with atopic dermatitis, it was found that S1P levels are decreased in lesional skin (Bäumer et al., 2011). However, in the Nc/Nga mouse model of dermatitis and the TNCB mouse model of dermatitis, the S1P Receptor modulator fingolimod, which downregulates activity of S1PRs 1, 3,4, and 5, was found to exert protective effects (Kohno et al., 2004; Sugita et al., 2010). In light of these studies, we asked whether S1P can act as an acute pruritogen in mice.

We set out to answer this question by examining the role of S1P in acute itch, and by dissecting the downstream molecular mechanisms in somatosensory neurons responsible for S1P-evoked behaviors and neuronal activation. Here we show that S1P can act as both a pruritogen and algogen. Our study is the first to identify a role for S1P signaling in acute itch, and to elucidate the downstream molecular mechanisms by which nociceptive and pruriceptive somatosensory neurons detect and respond to S1P. Our findings demonstrate the contribution of S1P signaling to cutaneous itch and pain, and have important applications for the design and use of S1PR modulators as therapeutics for chronic itch and pain diseases.

## Materials and Methods

### Mice

*S1pr3*^*mcherry/+*^ (Sanna et al., 2016) and *S1pr3* ^-/-^ (Kono et al., 2004) mice were obtained from Jackson Laboratory and backcrossed to C57bl6/J, *Trpv1*^-/-^ and *Trpa1*^-/-^ mice were described previously (Caterina et al., 2000; Bautista et al., 2006a) and *Trpv1*^-/-^/*Trpa1*^-/-^ mice were bred from crossing *Trpv1*^-/+^ and *Trpa1*^-/+^ mice. Mice (20–30 g; 8-10 weeks) were housed in 12 h light-dark cycle at 21°C. All experiments were performed under the policies and recommendations of the International Association for the Study of Pain and approved by the University of California, Berkeley Animal Care and Use Committee.

### Correlation of gene expression with itch

A previous study examined the correlation of transcript expression of individual genes in dorsal root ganglion (DRG) neurons with itch behavior among BXD mouse strains (Morita et al., 2015). One of the top correlated genes from the screen was *Fam57b* (*r* = 0.57), a recently identified ceramide synthase (Yamashita-Sugahara et al., 2013) and component of the S1P pathway that is robustly expressed in somatosensory neurons (Gerhold et al., 2013). To assess whether expression of the S1P pathway genes as a group was correlated with somatosensory behaviors across the mouse strains of the BXD population, we first tabulated the absolute value of the Pearson’s correlation *r* between expression of each S1P pathway gene in turn (*Fam57b, Ppargc1a, Spns1, Spns2, Sphk2, S1pr3, S1pr1, Esrrb, Esrrg, Lrp2*) and itch behavior (Morita et al., 2015), and calculated the median of these correlation values, *r*_*true*_. We then drew 10 random genes from the set of all 16,220 genes with detected expression and computed the median correlation as above using this null set, *r*_*null*_. Repeating the latter 10,000 times established a null distribution of median correlations; we took the proportion of resampled gene groups that exhibited (*r*_*true*_ ≥ *r*_*null*_) as an empirical p-value reporting the significance of enriched correlation between expression and itch in the genes of the S1P pathway.

### Mouse behavior

Itch and acute pain behavioral measurements were performed as previously described (Wilson et al., 2011). Mice were shaved one week prior to itch behavior. Compounds injected: 200 nM, 2 μM, 10 μM S1P (Tocris, Avanti Polar Lipids), in PBS with Methanol-PBS vehicle controls. Fresh S1P was resuspended in methanol and single-use aliquots were prepared and dried under nitrogen gas prior to use. Pruritogens were injected using both the neck/rostral back model (20 μl), and the cheek model (20 μl) of itch, as previously described (Wilson et al., 2011). Behavioral scoring was performed while blind to experimental condition and mouse genotype. All scratching and wiping behavior videos were recorded for 1 hour and scored for either the first 30 minutes (scratching) or the first five minutes (wiping). Bout number, time spent scratching, and bout length were recorded for scratching behavior. Wiping was recorded as number of wipes.

For radiant heat hypersensitivity behavior, S1P was injected intradermally into the plantar surface of the hindpaw (20 μl). Radiant heat paw withdrawal latencies, before and after application of compound or vehicle were performed as previously described (Tsunozaki et al., 2013; Morita et al., 2015) using the Hargreaves test system (IITC Life Science). Mice were injected with compound of interest into the hind paw, paw withdrawal latencies were measured 15 min pre- and 20 min-30 min post-injection. Heat-evoked responses included fast paw withdrawal, licking/biting/shaking of the affected paw, or flinching. Mice were allowed to acclimate on platform for 1 hour before injection. The radiant heat source raised the platform temperature to 39.8 °C within 5 seconds, and to 60 °C within 10 seconds, as measured by a fast temperature probe (Physitemp).

Wherever possible, wild-type littermate controls were used in behavioral experiments. Mice were singly housed one week prior to all behavioral experiments. All mice were acclimated in behavioral chambers on the 2 subsequent days for at least 1 hour prior to treatment for itch/pain behavior and radiant heat. Age-matched or littermate male mice were used for all behavioral studies. Mice were tested in 4-part behavior chambers (IITC Life Science) with opaque dividers (Tap Plastics). Itch and acute pain behavior was filmed from below using high-definition cameras.

### In situ hybridization (ISH)

ISH was performed as previously described. Fresh DRG were dissected from 8-12 week old mice, flash frozen in OCT embedding medium, and sectioned at 14 μm onto slides. ISH was performed using Affymetrix Quantigene ViewISH Tissue 2-plex kit according to manufacturer’s instructions with Type 1 and Type 6 probes. The following probes against mouse mRNAs were created by Affymetrix and used for ISH: *S1pr3* and *Mrgpra3.* Slides were mounted in Fluoromount with No. 1.5 coverglass. Imaging of ISH experiments and all other live- and fixed-cell imaging was performed on an Olympus IX71 microscope with a Lambda LS-xl light source (Sutter Instruments). Images were analyzed using ImageJ software. Briefly, DAPI-positive cells were circled and their fluorescence intensity (AFU) for all channels was plotted against cell size using IgorPro software. Co-labeling analysis was performed using ImageJ. Intensity thresholds were set based on the negative control staining slide. Cells were defined as “co-expressors” if their maximum intensities were greater than the threshold for both the Type 1 and Type 6 probe channels.

### Cell culture

Cell culture was carried out as previously described (Wilson et al., 2011). Briefly, neurons from dorsal root ganglia of 2-8 week old male and female mice, or trigeminal ganglia from P0-P4 neonates, where indicated, were dissected and incubated for 10 min in 1.4 mg ml−1 Collagenase P (Roche) in Hanks calcium-free balanced salt solution, followed by incubation in 0.25% standard trypsin (vol/vol) -EDTA solution for 2 min with gentle agitation. Cells were then triturated in media (MEM Eagle’s with Earle’s BSS medium, supplemented with 10% horse serum (vol/vol), MEM vitamins, penicillin/streptomycin and L-glutamine), plated onto Poly D-Lysine (Sigma, 1 mg/mL) and Laminin (Corning, 1:300) coated glass coverslips and used within 20 h.

### Calcium imaging

Ca^2+^ imaging experiments were carried out as previously described (Wilson et al., 2011). DRG neurons from 2-8 week old male and female mice or P0-4 TG from mice of unknown sex were used for all experiments. An age-matched wild-type control was also prepared and imaged the same day for experiments on knockout neurons. Cells were loaded for 60 min at room temperature with 10 μM Fura-2AM supplemented with 0.01% Pluronic F-127 (wt/vol, Life Technologies) in a physiological Ringer’s solution containing (in mM) 140 NaCl, 5 KCl, 10 HEPES, 2 CaCl2, 2 MgCl2 and 10 D-(+)-glucose, pH 7.4. All compounds were purchased from Sigma. Acquired images were displayed as the ratio of 340 nm/380 nm. Cells were identified as neurons by eliciting depolarization with high potassium Ringer’s solution (75 mM) at the end of each experiment. Responding neurons were defined as those having a > 15% increase from baseline ratio. Image analysis and statistics were performed using automated routines in Igor Pro (WaveMetrics). Fura-2 ratios were normalized to the baseline ratio F340/F380 = (Ratio)/(Ratio t = 0).

### Electrophysiology

Current clamp experiments were carried out as previously described (Hill et al., 2018). Briefly, gap-free current clamp recordings were collected at 5 kHz and filtered at 2 kHz (Axopatch 200B, pClamp software). Electrode resistance ranged between 2–5 MΩ. Internal solution contained 140 mM KCl, 2 mM MgCl2, 1 mM EGTA, 5 mM HEPES, 1 mM Na2ATP, 100 μM GTP, and 100 μM cAMP (pH 7.4). Bath solution was physiological Ringer’s solution. The pipette potential was canceled before seal formation. Experiments were carried out only on cells with a series resistance of less than 30 MΩ and membrane capacitance of < 40 pF. Current injection was used to stabilize cells to ~ −60 mV prior to experiment. Action potentials which occurred +/- 1 sec of drug addition or a gravity perfusion artifact were not counted as responses to the drug. For experiments where two drugs were added in succession, an increase in spike frequency was considered a response for the second drug. For recordings from *S1pr3*^*mCherry/+*^ animals, S1PR3+ DRG neurons were identified using a standard fluorescence microscope. Analysis of electrophysiology data was performed in pClamp and IgorPro. Experimenter was blinded to the antagonists applied until after data analysis was completed.

### Experimental design & statistical analyses

All statistical tests were performed using Prism (GraphPad). Values are reported as the mean ± SEM (where N = number of mice used) for calcium imaging experiments where multiple independent days of imaging were performed, and mean ± SD for all other experiments (N = number of wells or number of mice). A one- or two-way ANOVA followed by the Sidak’s, Dunnett’s or Tukey’s post hoc tests (where appropriate) was employed. Number of mice or samples required to attain significance was not calculated beforehand and was based on numbers used in similar behavioral studies. For behavioral experiments, mice were randomly assigned to treatment groups by the individual who blinded the experimenter. For behavioral experiments, every effort was made to ensure equal numbers of mice of each genotype were used for each experiment (where appropriate), and that treated and control groups were of identical or near-identical size. Significance was labeled as: n.s., not significant, p ≥ 0.05; *p < 0.05; **p < 0.01; ***p < 0.001.

## Results

### S1P triggers itch via S1PR3

Our group previously harnessed natural variation in somatosensory behaviors among genetically distinct mouse strains to identify candidate transducers in dorsal root ganglion (DRG) somatosensory neurons (Morita et al., 2015). Analysis of this dataset revealed that members of the sphingosine 1-phosphate (S1P) synthesis and signaling pathways (Fig. 1a) were expressed in somatosensory neurons in a manner that correlated with itch behaviors (Fig. 1b). Previous studies have shown that S1P promotes acute pain (Camprubi-Robles et al., 2013) and thermal sensitization via S1P Receptor 3 (S1PR3), and is required for normal mechanical pain sensitivity (Hill et al., 2018). Here, we sought to address the specific contribution of S1P/S1PR3 signaling to S1P-evoked itch behaviors.

**Figure 1.**
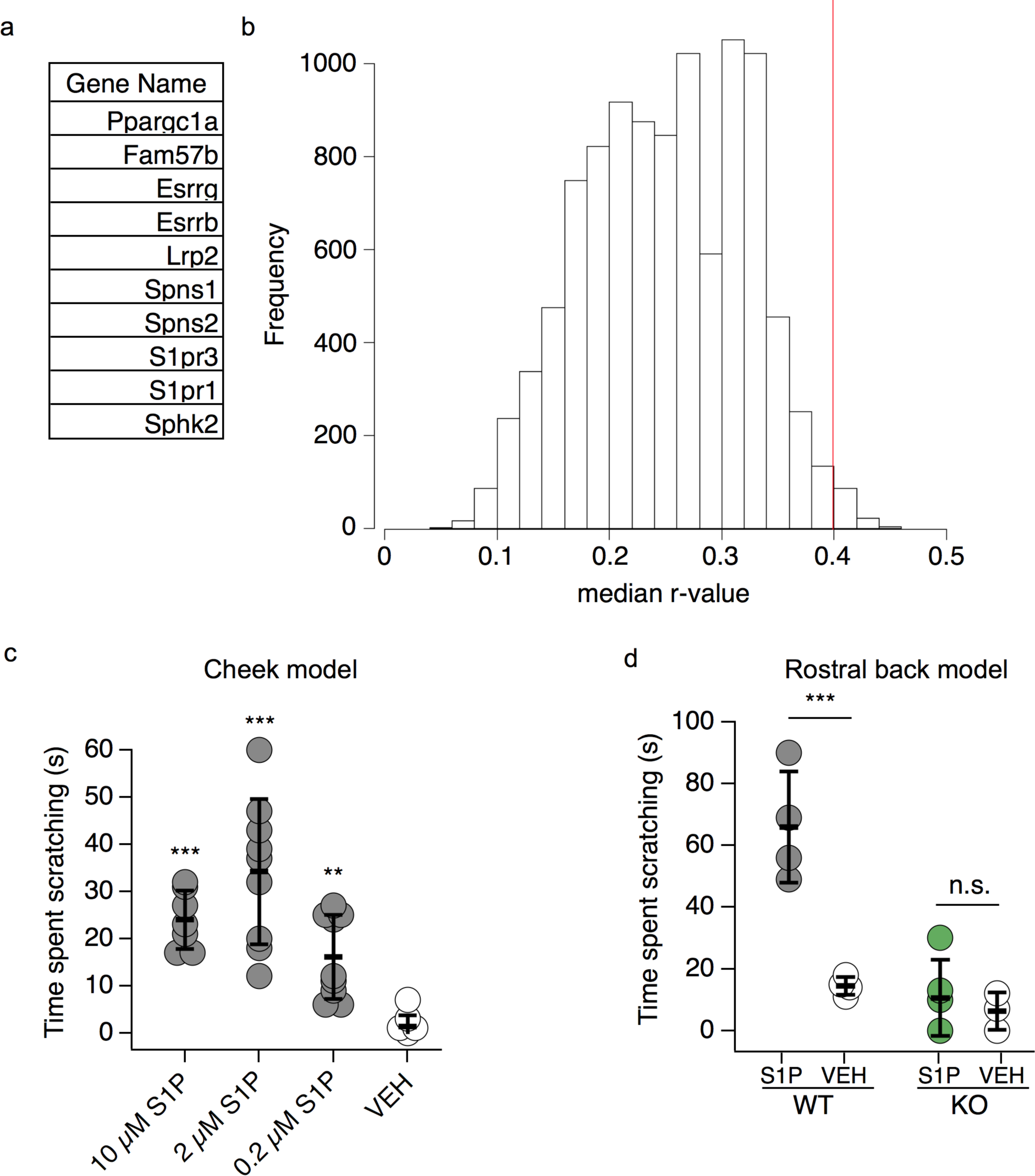
S1P triggers itch via S1PR3. **a.** Tabular list of the top 10 most highly expressed sphingosine 1-phosphate (S1P) pathway genes in dorsal root ganglion (DRG) neurons. **b.** Histogram showing median absolute r-value for gene expression vs. itch plotted against frequency for 10,000 permutations of random subsets of 10 genes in the BXD DRG transcriptome. Red line indicates median r-value for S1P genes (*p* = 0.0116, r_median_ = 0.399, see Methods). **c.** (Left) Intradermal cheek injection of 10 μM S1P, 2 μM, 0.2 μM, and 20 μL 1% methanol PBS (VEH), with quantification of time spent scratching over the 30-minute post-injection interval; *p* < 0.0001 (one-way ANOVA (F(3,30) = 18.29); N = 7-9 mice per condition). Dunnett’s multiple comparisons *p*-values are represented on graph for comparisons made between treated and vehicle groups. **d.** Time spent scratching in response to intradermal injection of 20 μL 0.2 μM S1P or 0.1% methanol-PBS vehicle in rostral back of age-matched *S1pr3* ^+/+^ (WT) and *S1pr3* ^-/-^ (KO) mice; *p* = 0.0089 (one-way ANOVA (F(3,9) = 7.274); N = 3-4 mice per treatment). Tukey’s multiple comparisons *p*-values are represented on graph for comparisons made between treated and vehicle groups. Error bars represent mean ± SD.

We first asked whether exogenous S1P can trigger itch using the cheek assay, which allows for simultaneous discrimination of pain-evoked wiping and itch-evoked scratching in mice (Shimada and LaMotte, 2008). Injection of 2-10 μM S1P triggered acute nocifensive behaviors, as previosuly shown (Camprubi-Robles et al., 2013; Hill et al., 2018). However, we also observed robust scratching in these animals (Fig. 1c). Intriguingly, 200 nM S1P elicited robust scratching behaviors (Fig. 1c), but no pain behaviors. Scratching behaviors developed with an average latency of 4 minutes 29 seconds ± 1 minute 37 seconds and persisted for at least 30 minutes. We also observed that injection of 200 nM S1P into the rostral back triggered itch behaviors with an average scratching time of 66 ± 18.1 seconds, which was significantly greater than scratching evoked by vehicle (0.1% methanol-PBS; 14.5 ± 2.88 seconds; Fig. 1d). To begin to elucidate the mechanisms underlying these effects, we focused on the S1P receptor S1PR3, which is required for S1P-evoked pain and pain hypersensitivity (Camprubi-Robles et al., 2013; Hill et al., 2018). Assessment of itch behavior in *S1pr3*^-/-^ mice revealed significantly attenuated S1P-evoked scratching, indistinguishable from vehicle injection (10.6 ± 12.3 s versus 6.3 ± 6.2 s, respectively; Fig. 1d). Furthermore, in a previous report, we observed no defects in itch responses to the pruritogens chloroquine and histamine in *S1pr3*^-/-^ mice (Hill et al., 2018). These data suggest that S1PR3 is the primary receptor by which S1P signals in somatosensory neurons to drive itch. Our findings show that S1P can act as a pruritogen, selectively triggering itch at nanomolar concentrations, whereas at micromolar concentrations, S1P acts both as a pruritogen and algogen, triggering itch and pain. Our discovery that S1PR3 is required for S1P-evoked itch is consistent with previous studies showing that S1P-evoked acute pain (Camprubi-Robles et al., 2013), heat hypersensitivity, and mechanical pain (Hill et al., 2018) are absent in *S1pr3*^-/-^ mice. These results add S1P to a growing list of endogenous molecules that act as both pruritogens and algogens (Shimada and LaMotte, 2008; Akiyama and Carstens, 2013; Moore et al., 2017; Esancy et al., 2018).

### S1PR3 is functional and expressed in a subset of pruriceptors

We next asked whether S1PR3 is functional and expressed in itch neurons. Calcium imaging of S1P responses in wild-type mouse somatosensory neurons revealed that 1 μM S1P activates 34.2% of all neurons, which fall into two main populations: 22.6% (66.1% of S1P^+^ neurons) are TRPV1^+^/TRPA1^+^ neurons, which are capsaicin- and allyl isothiocyanate (AITC)-sensitive, and 11.6% (33.9% of S1P^+^ neurons) are TRPV1^+^/TRPA1^-^ neurons, which are capsaicin-sensitive and AITC-insensitive (Fig. 2a,b). Neurons which were responsive to both AITC and S1P exhibited an EC_50_ for S1P of 102 nM, and were activated by lower S1P concentrations (10 nM) than neurons which were responsive to capsaicin and S1P (EC_50_ = 155 nM) and showed robust responses to 100 nM S1P (Fig. 2c). The complete overlap of S1P-responsive neurons with TRPV1^+^ neurons is consistent with a role for S1P in itch and pain. Previous studies have shown that the MrgprA3+ subpopulation of TRPA1^+^ neurons is required for many forms of non-histaminergic acute and chronic itch (Liu et al., 2009, 2012; Wilson et al., 2011; Han et al., 2013; Reddy et al., 2015; Zhu et al., 2017). In keeping with our hypothesized role for S1P in itch, 80% of chloroquine-responsive MrgprA3^+^ neurons responded to S1P (Fig. 2b). We also observed a population of S1P-sensitive neurons which were sensitive to the pruritogen histamine (23.6% of S1P-sensitive neurons; Fig. 2b). We conclude that S1P activates thermal nociceptors that express TRPV1 and pruriceptors that express TRPA1 and TRPV1.

**Figure 2.**
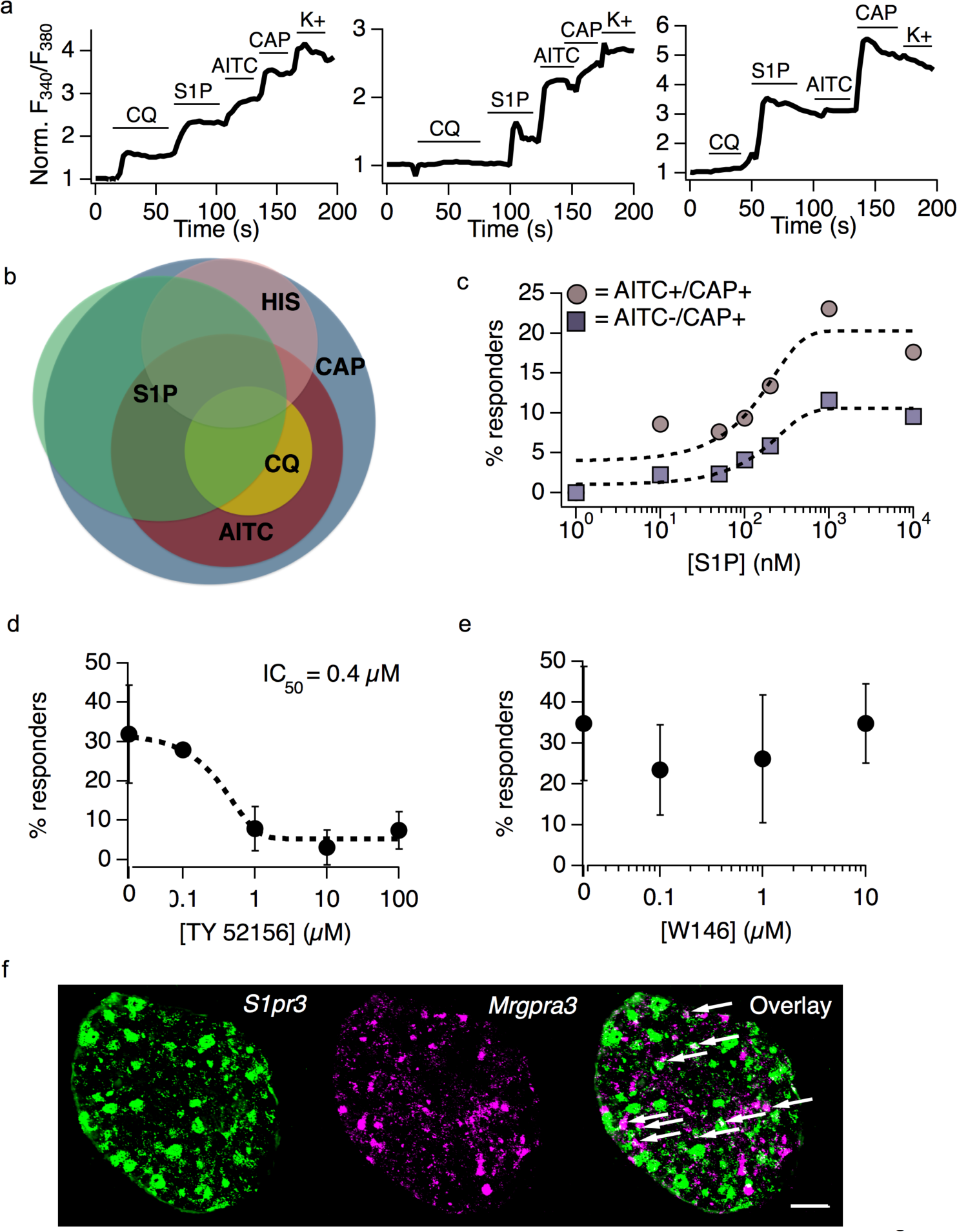
S1PR3 is functional and expressed in a subset of pruriceptors. **a.** Representative calcium imaging traces showing normalized Fura-2 signal (Norm. F340/F380) over time before and after addition of indicated compounds in cultured DRG neurons (CQ = 1 mM chloroquine, S1P = 1 μM S1P; AITC = 100 μM allyl isothiocyanate; CAP = 1 μM capsaicin; K+ = Ringer’s with 75 mM potassium). (Left) Putative pruriceptor that responds to both chloroquine and S1P; (Center) putative pruriceptor or nociceptor which responds to both AITC and S1P; (Right) Putative thermal nociceptor which responds to S1P and capsaicin but not AITC. **b.** Venn diagram demonstrating overlap of neuronal subpopulations activated by capsaicin, AITC, histamine (HIS; 100 μM) chloroquine, and S1P. Relative proportion of overlap represents % overlap as calculated from ratiometric calcium imaging data (N = 4,000 neurons). **c.** Dose-response curve showing proportion of DRG neurons responding to varying concentrations of S1P. Concentrations used: 1, 10, 50, 100, 200, 1000, and 10,000 nanomolar (N = 2 animals). Colored traces indicate proportion of neurons responding to S1P and AITC or S1P and capsaicin at indicated concentrations. **d.** S1P-evoked calcium transients in DRG neurons are inhibited by the selective S1PR3 antagonist TY 52156. Black dotted line indicated sigmoidal fit from which IC_50_ was derived (N = 2 animals). Error bars represent mean ± SD. **f.** S1P-evoked calcium transients in DRG neurons are not inhibited by the selective S1PR1 antagonist W146 (N = 2 animals). Error bars represent mean ± SD. **f.** Co-ISH of *S1pr3* (green) with *Mrgpra3* (magenta) in sectioned whole DRG from adult wild-type mice. Third column: overlay. Scale bar = 100 μm. Images were acquired using a 10x air objective. Arrows indicate cells showing co-expression of both markers.

To pursue further the mechanism by which S1P acts in somatosensation, we examined the effects of pharmacological blockade of S1PR3 and S1PR1 on S1P-evoked calcium responses. Incubating cells with the S1PR3-selective antagonist TY 52156 attenuated S1P-evoked calcium transients with an IC_50_ of 0.4 μM (Fig. 2d). In contrast, the S1PR1-selective antagonist W146 had no discernable effect on S1P responses at a range of concentrations that exceed reported inhibitory concentrations (70-80 nM) for this drug (Fig. 2e; Finley et al., 2013; Janes et al., 2014). These data support previous studies showing that cultured somatosensory neurons from *S1pr3*^-/-^ mice exhibit no calcium responses to 1 μM S1P (Camprubi-Robles et al., 2013; Hill et al., 2018) and suggest that S1PR3, but not S1PR1, is required for activation of pruriceptors and nociceptors by S1P.

While S1PR3 is expressed (Hill et al., 2018) and functional in distinct subsets of TRPV1^*+*^ and TRPV1+/TRPA1^*+*^ DRG neurons, whether this receptor is expressed in pruriceptors was unknown. To investigate this, we performed co-*in situ* hybridization (co-ISH) of *S1pr3* with *Mrgpra3*, which encodes the chloroquine receptor that marks a neuronal subpopulation required for some forms of acute and chronic itch (Liu et al., 2009). We found that 35.1% of *Mrgpra3*^*+*^ cells express *S1pr3* and 11.6% of *S1pr3*^+^ cells express *Mrgpra3* (Fig. 2f). The differing proportions of *S1pr3*^*+*^/*Mrgpra3*^*+*^ neurons in our ISH and of S1P^+^/chloroquine^+^ neurons in our calcium imaging studies (Fig. 2b) may reflect a difference between intact DRG versus cultured DRG neurons, and/or the sensitivity of mRNA expression versus calcium imaging assays. In summary, our calcium imaging and co-ISH data, along with our previous findings, reveal two main populations of small-diameter *S1pr3*^*+*^ neurons: 1) *Trpv1*^+^ thermal nociceptors and, 2) *Trpa1*^+^ cells, including a subset of *Mrgpra3*^*+*^ pruriceptors. These results dovetail with our finding that S1PR3 is required for S1P-evoked itch (Fig. 1d).

### S1P triggers neuronal activation in distinct subsets of nociceptors and pruriceptors

We next sought to identify components downstream of S1PR3 underlying S1P-evoked calcium transients in cultured mouse somatosensory neurons. S1P responses were mediated solely by calcium influx, as chelation of extracellular calcium abolished all responses (Fig. 3a), suggesting that S1P triggers the opening of plasma membrane cation channels. We also observed similar results with application of Ruthenium Red (Fig. 3b), a blocker of a number of calcium-permeable ion channels, including Transient Receptor Potential (TRP) channels.

**Figure 3.**
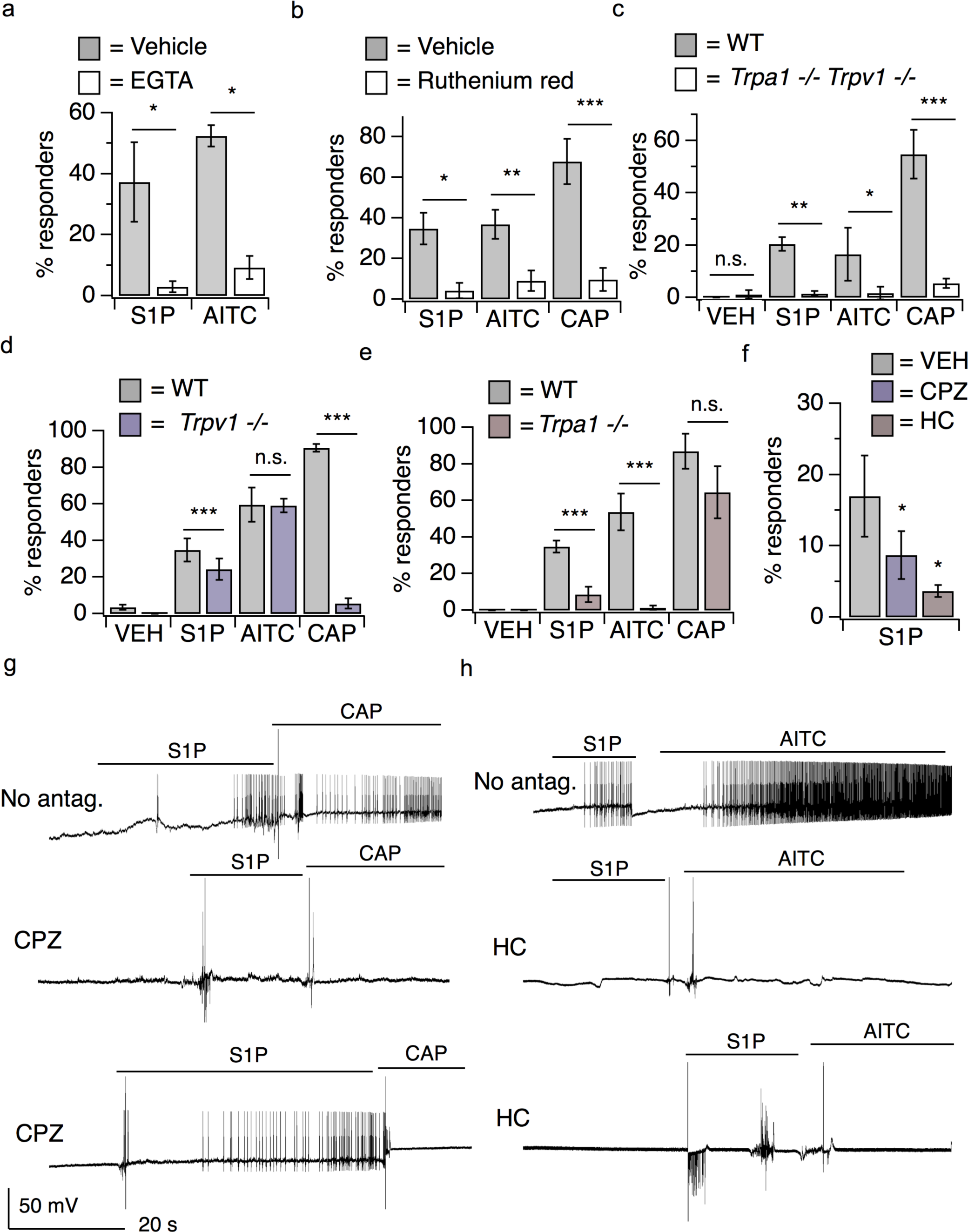
S1P triggers neuronal activation in distinct subsets of nociceptors and pruriceptors. **a.** Percent DRG neurons responding to S1P and AITC in Calcium Ringer’s and 10 μM EGTA Ringer’s; *p* < 0.0001 (one-way ANOVA (F(3,11) = 8.291); N = 4 wells of ~100 neurons each per treatment). Error bars represent mean ± SD. **b.** Percent responders to 1 μM S1P, AITC, and Cap in vehicle and 10 μM Ruthenium Red; *p* < 0.0001 (one-way ANOVA (F(5,17) = 17.42); N = 5 wells of ~100 neurons each per treatment). Error bars represent mean ± SD. **c.** Percent responders to vehicle, S1P, AITC, and capsaicin in *Trpa1/Trpv1* ^+/+^ and ^-/-^ DRG neurons; *p* < 0.0001 (one-way ANOVA (F(7,40) = 8.555); N = 2 age-matched mice per genotype) Error bars represent mean ± SD. **d.** Percent responders to vehicle, S1P, AITC, and capsaicin in *Trpv1 +/+* and *Trpv1* -/- neurons; *p* < 0.0001 (one-way ANOVA (F(7, 179) = 72.67); N = 3 age-matched mice per genotype). Error bars represent mean responses ± SEM. **e.** Percent responders to vehicle, S1P, AITC, and capsaicin in *Trpa1 +/+* and *Trpa1* -/- neurons; *p* < 0.0001 (one-way ANOVA (F(7,210) = 105); N = 3 age-matched mice per genotype). Error bars represent mean responses ± SEM. **f.** Effect of 50 μM Capsazepine (TRPV1 antagonist) and 50 μM HC-030031 (TRPA1 antagonist) vs. DMSO-Ringer’s vehicle on S1P-evoked calcium responses in cultured DRG neurons; *p* < 0.0001 (one-way ANOVA (F(8,30) = 84.32); N = 3 wells of ~100 neurons each per treatment) Error bars represent mean responses ± SD. Unless otherwise indicated, Sidak’s multiple comparisons were made between treated and vehicle for each agonist. **g.** Whole cell current-clamp recordings of S1PR3+ DRG neurons from *S1pr3*^*mCherry/+*^ animals pre-treated for 5-10 minutes with Vehicle (1% DMSO in Ringer’s) or 50 μM Capsazepine and subsequently exposed to 1 μM S1P followed by 1 μM capsaicin, indicated by lines. (Top) Recording from a neuron that responded to S1P and CAP; (Middle) A neuron that responded to neither S1P nor CAP; (Bottom) a neuron that responded to S1P but not CAP. In these experiments, 5 of 6 S1PR3+ cells which received vehicle responded to S1P and 5 of 6 responded to CAP; 3 of 8 cells which received Capsazepine responded to S1P and 0 of 8 cells responded to CAP (N = 3 animals). **h.** Whole cell current-clamp recordings of S1PR3+ DRG neurons from *S1pr3*^*mCherry/+*^ animals pre-treated for 5-10 minutes with Vehicle (1% DMSO in Ringer’s) or 50 μM HC-030031 and subsequently exposed to 1 μM S1P followed by 100 μM AITC, indicated by lines. (Top) Recording from a neuron that responded to S1P and AITC; (Middle) A neuron that responded to neither S1P nor AITC; (Bottom) a neuron that responded to S1P but not AITC. In these experiments, 4 of 5 S1PR3+ cells which received vehicle responded to S1P and 2 of 5 responded to AITC; 4 of 11 cells which received HC-030031 responded to S1P and 0 of 11 cells responded to AITC (N = 3 animals).

Since we observed that all S1P responsive neurons expressed TRPA1 and/or TRPV1, and these channels have been shown to functionally couple to a variety of GPCRs (Chuang et al., 2001; Prescott and Julius, 2003; Bandell et al., 2004; Dai et al., 2007; Shim et al., 2007; Kwon et al., 2008; Rohacs et al., 2008; Imamachi et al., 2009; Schmidt et al., 2009; Wilson et al., 2011; Moore et al., 2017), we asked whether either channel mediated S1P-evoked neuronal activation. Neurons isolated from mice lacking both TRPA1 and TRPV1 exhibited greatly attenuated S1P-evoked calcium responses, similar to those evoked by vehicle (Fig. 3c). Intriguingly, we found the percentage of S1P-responsive neurons was partially attenuated in neurons from *Trpv1*^*-/-*^ (Fig. 3d) and *Trpa1*^-/-^ single knockout mice (Fig. 3e). Such genetic effects were mirrored by pharmocological blockade of these channels: the TRPV1 antagonist capsazepine (50 μM; CPZ) or the TRPA1 antagonist HC-030031 (50 μM; HC) were sufficient to fully block capsaicin or AITC responses, respectively, and resulted in a significant, though partial, attenuation of neuronal S1P responses (Fig. 3f).

To test whether TRPA1 and/or TRPV1 are required for S1P-evoked neuronal excitability, we turned to current-clamp whole cell electrophysiology to examine action potential (AP) firing. For this purpose, we isolated and cultured DRG neurons from the *S1pr3*^mCherry^ reporter mouse, which produces a functional S1PR3-mCherry fusion protein (Sanna et al., 2016), and examined their changes in membrane potential in response to S1P. Our experiments revealed that 1 μM S1P elicited AP firing in both the capsaicin-sensitive and AITC-sensitive S1PR3+ populations of sensory neurons (Fig. 3g-h, top). We next used a pharmacological approach to investigate the role of TRPV1 and TRPA1 in S1P-evoked AP firing. TRPV1 blockade with the antagonist CPZ decreased the proportion of neurons which fired APs in response to S1P and, as expected, completely blocked capsaicin-evoked AP firing in these neurons (Fig. 3g). Likewise, blockade of TRPA1 with HC resulted in a decreased proportion of cells which fired in response to S1P and the expected complete loss of AITC-evoked firing (Fig. 3h). These results suggest that TRPV1 and TRPA1 directly contribute to S1P-evoked neuronal activation and AP firing. Combined with our calcium imaging studies using TRP channel inhibitors and A1/V1 knockout neurons, our data show that TRPA1 and TRPV1 are required for S1P-evoked neuronal activation in distinct populations of somatosensory neurons.

**S1PR3 utilizes distinct G-protein coupled pathways to activate subsets of somatosensory neurons** We used a pharmacological approach to investigate the mechanisms by which S1PR3 signals to TRPA1 and TRPV1. We first focused on phospholipase C (PLC), a known target of the S1PR3 signaling partner G_q_ (Kim et al., 2011; Flock et al., 2017). Given that GPCRs have been shown to activate TRPV1 and TRPA1 channels via PLC activity in somatosensory neurons (Chuang et al., 2001; Prescott and Julius, 2003; Bandell et al., 2004; Shim et al., 2007; Dai et al., 2007; Kwon et al., 2008; Rohacs et al., 2008; Schmidt et al., 2009; Imamachi et al., 2009; Wilson et al., 2011; Paulsen et al., 2015; Gao et al., 2016; Moore et al., 2017), we hypothesized that PLC signaling could be required for S1P-evoked calcium responses in sensory neurons. This notion bore out, in that inhibition of PLC signaling using the drug U73122 significantly decreased the percentage of neurons that displayed S1P-evoked calcium responses (Fig. 4a). This effect was most pronounced in the TRPA1^-^ population of S1P-responsive neurons (Fig. 4b,c), but had only a small effect on the TRPA1^+^ population (Fig. 4b,c). Next, given that GPCR signaling via G_βγ_ can activate TRPA1 (Wilson et al., 2011), we asked if G_βγ_ signaling could also play a role in S1P-evoked calcium responses. We observed a significant reduction in the percentage of neurons responding to S1P upon blockade of G_βγ_ activity using the drug gallein (Fig. 4a). And in contrast to PLC inhibition, gallein’s effects were more robust in the TRPA1^+^ population and were minimal in the TRPA1^-^ population (Fig. 4b,c). Blockade of both pathways using U73122 and gallein resulted in a significant loss of all neurons responsive to S1P (Fig. 4a), irrespective of population (Fig. 4b,c). Thus, S1PR3 can signal in two different sensory neuronal subtypes (TRPA1^+^/TRPV1^+^ and TRPA1^-^/TRPV1^+^ neurons) in a concentration-dependent manner, using distinct molecular signaling molecules (G_βγ_ and PLC, respectively).

**Figure 4.**
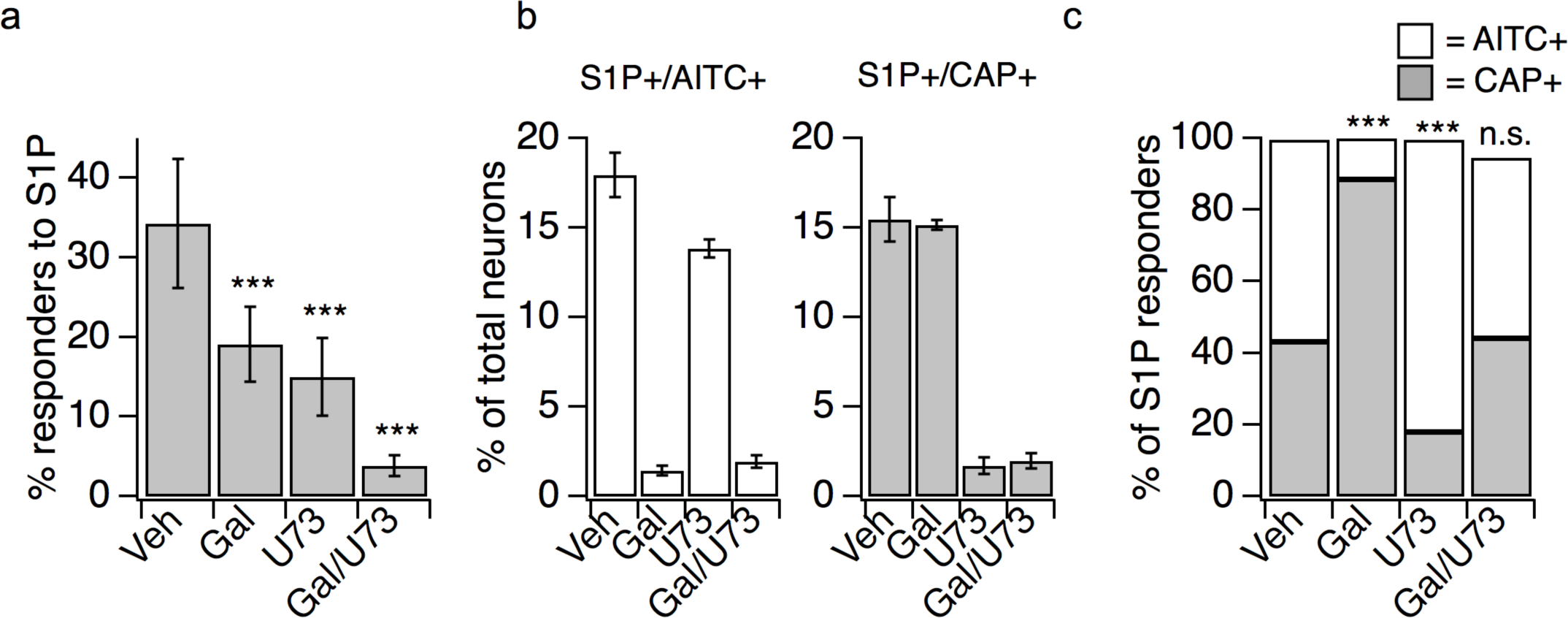
S1PR3 utilizes distinct G-protein-coupled pathways to activate subsets of somatosensory neurons. **a.** Percent of DRG neurons responding to 1 μM S1P after 15 minute incubation with either DMSO-Ringer’s vehicle (Veh), 100 μM Gallein (Gal; GβΓ blocker), 1 μM U73122 (U73; PLC blocker), or U73122 + Gallein; *p* < 0.0001 (one-way ANOVA (F(3,28) = 31.46); N = 4-10 wells of ~50 neurons each per treatment from 2 animals). Error bars represent mean responses ± SD. Dunnett’s multiple comparison *p-*values are indicated on graph for comparisons to vehicle. **b.** From experiments in **(a)**, percent of total neurons which were sensitive to S1P, AITC and capsaicin (white, left) or S1P and capsaicin only (grey, right); Error bars represent mean responses ± SD. **c.** From experiments in **(a),** percentage of S1P-responsive neurons that were sensitive to AITC and capsaicin (white) or capsaicin only (grey); *p* < 0.0001 (one-way ANOVA (F(3,28) = 41.48). Error bars are ommitted for clarity, as they are represented in **(b),** prior to normalization to percent of S1P-responsive neurons. Dunnett’s multiple comparison *p-*values are indicated on graph for comparisons to vehicle.

### S1P evokes itch and pain behaviors via distinct TRP channels

Here we have shown that TRPA1 and TRPV1 are required for S1P-evoked neuronal exicitation and calcium responses in pruriceptors and nociceptors. We thus tested the requirement of TRPA1 and TRPV1 to S1P-evoked itch and pain behaviors. We observed that S1P evoked robust heat hypersensitivity in wild-type and *Trpa1*^-/-^ mice, but not in *Trpv1*^-/-^ mice (Fig. 5a). Similarly, 10 μM S1P triggered acute nocifensive behaviors (wiping) in wild-type and *Trpa1*^-/-^ mice, but not in *Trpv1*^-/-^ mice (Fig. 5b). Finally, we examined S1P-evoked itch behaviors in wild-type, *Trpa1*^-/-^ and *Trpv1*^-/-^ mice using the rostral back model. In contrast to our pain data, 200 nM S1P evoked robust scratching behaviors in wild-type and *Trpv1*^-/-^ mice, but not in *Trpa1*^-/-^ mice (Fig. 5c). These data support a model whereby TRPV1 selectively mediates S1P-evoked acute pain and heat hypersensitivity, whereas TRPA1 selectively mediates S1P-evoked acute itch (Figure 6).

**Figure 5.**
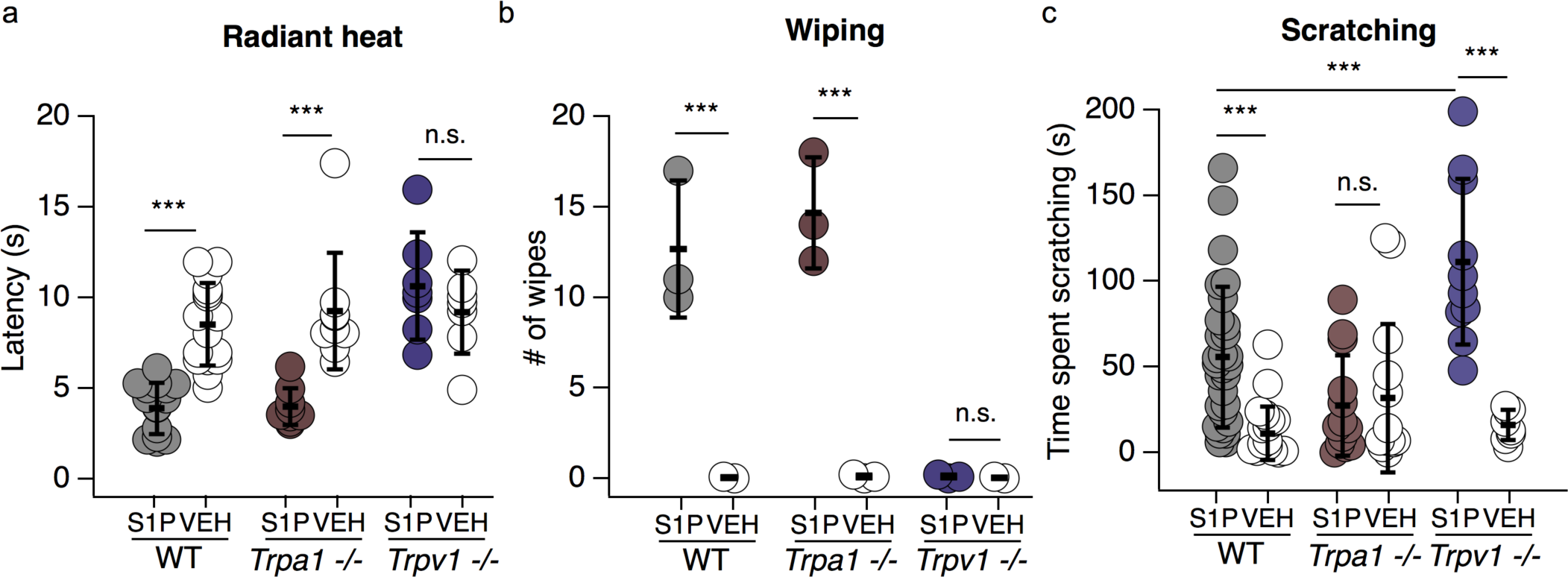
S1P evokes itch and pain via distinct TRP channel-dependent pathways. **a.** Radiant heat paw withdrawal latencies 20-30 minutes post injection of 15 μL 10 μM S1P or 0.3% methanol-PBS vehicle i.d. into the hind paw of age matched wild-type, *Trpa1* ^-/-^ and *Trpv1* ^-/-^ mice; *p* < 0.0001 (one-way ANOVA (F(5,66) = 14.13); N = 7 or more age-matched mice per condition). **b.** Number of wipes in response to intradermal (i.d.) injection of 20 μL 10 μM S1P or 0.3% methanol-PBS vehicle in cheek of age-matched wild-type, *Trpa1* ^-/-^ and *Trpv1* ^-/-^ mice; *p* < 0.0001 (one-way ANOVA (F(5,11) = 33.98); N = 3 age-matched mice per condition). **c.** Time spent scratching in response to intradermal injection of 20 μL 0.2 μM S1P or 0.1% methanol-PBS vehicle in rostral back of age-matched wild-type, *Trpa1* ^-/-^ and *Trpv1* ^-/-^ mice; *p* < 0.0001 (one-way ANOVA (F(5,91) = 14.13); N = 8 or more age-matched mice per condition). For all graphs, Sidak’s multiple comparisons were made between all genotypes and treatments and error bars represent mean ± SD.

**Figure 6.**
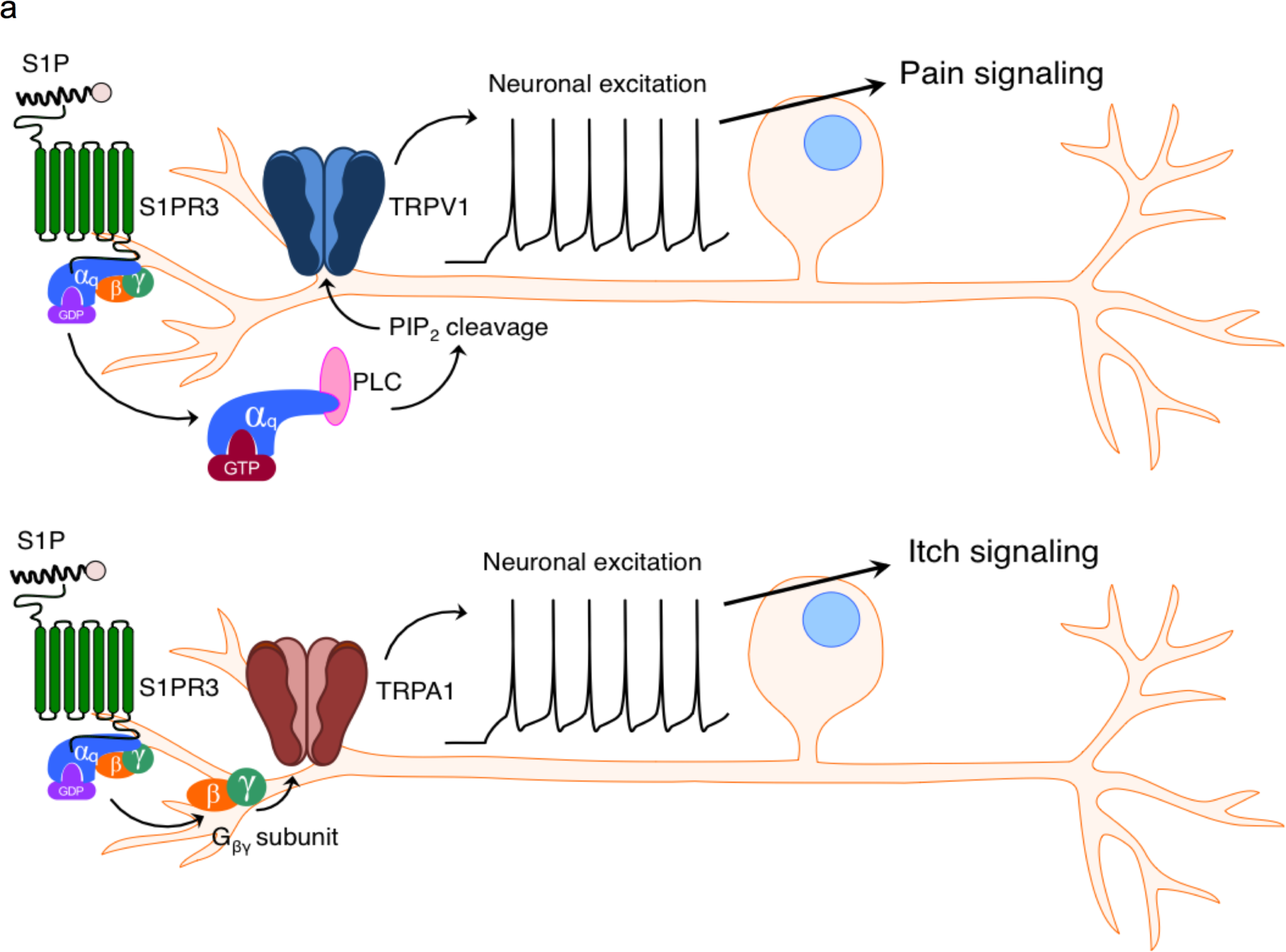
The effects of S1P on thermal hypersensitivity and acute pain and itch. **a.** Elevated S1P 1) elicits thermal hypersensitivity and acute pain via S1PR3-dependent activation of TRPV1, which requires PLC activity (top) and 2) elicits itch-evoked scratching via S1PR3-dependent activation of TRPA1, which requires G_βΓ_ (bottom).

## Discussion

Recent studies have linked aberrant S1P signaling to a variety of diseases, including: asthma (Chiba et al., 2010; Trankner et al., 2014; Roviezzo et al., 2015), multiple sclerosis (Brinkmann et al., 2010; Choi et al., 2011), cancer (Liang et al., 2013), psoriasis (Checa et al., 2015; Myśliwiec et al., 2016), and chronic pain (Patti et al., 2008; Janes et al., 2014; Grenald et al., 2017). Yet the cellular and molecular targets of S1P in these disorders are largely unknown. Here, we show that S1PR3 is required for S1P-evoked pain and itch, highlighting the potential for a direct contribution of S1P to sensitization, pain and/or itch in these diseases.

There is much interest in exploring the therapeutic potential of S1PR signaling in chronic pain and itch. However, little was known about the role of S1PRs in sensory neurons and their role in itch. We now show that S1P activates pruriceptors and triggers itch via S1PR3 and TRPA1. In mice, we found that injection of μM S1P triggered acute itch behaviors while concentrations at or above 2 μM triggered both itch and pain. S1PR3-deficient mice displayed a complete loss of S1P-evoked acute itch (Fig. 1d) and pain (Hill et al., 2018).

Our findings suggest sensory neuronal S1PR3 may play roles in disorders already linked to S1P signaling. Although S1P is produced by nearly every cell type, blood plasma contains the highest levels of S1P (low micromolar), and S1P is secreted in large amounts by activated mast cells (Idzko et al., 2002; Rivera et al., 2008; Olivera et al., 2013; Roviezzo et al., 2015; Saluja et al., 2017). Others have shown that sensory neurons innervating the lung express *S1pr3* and that administration of a non-selective S1P receptor agonist can enhance asthma-like airway hyperreactivity after acetylcholine challenge (Trankner et al., 2014), and that sensory neurons of the nodose ganglia are directly activated by S1P (Wang et al., 2017). It is also well known that TRPV1+ sensory afferents are required for airway hypersensitivity (Rogerio et al., 2011). Furthermore, mast cells, which release S1P when activated (Saluja et al., 2017), play important roles in allergic asthma as well as itch, and we propose that S1PR3 may play an important role in mediating itch and/or airway hyperreactivity under these conditions.

We have identified two distinct populations of sensory neurons which are excited by S1P: the TRPA1^+^/TRPV1^+^ and TRPA1^-^/TRPV1^+^ populations, that have been shown to differentially contribute to itch and pain behaviors (Caterina et al., 2000; Bandell et al., 2004; Bautista et al., 2006b; Dai et al., 2007; Shim et al., 2007; Gerhold and Bautista, 2009; Imamachi et al., 2009; Wilson et al., 2011; Moore et al., 2017). Our observations that the TRPA1^+^ population is activated by lower doses of S1P than the TRPA1^-^ population (Fig. 2c), and that itch behaviors are triggered by doses which are non-painful (Fig. 1c), supports a recent study showing that both zebrafish and mouse pruriceptors are significantly more sensitive to stimuli than nociceptors. This increased sensitivity results in selective recruitment of pruriceptors over nociceptors by the same agonist in a dose-dependent manner, such that lower concentrations of agonist selectively trigger itch and higher concentrations trigger both itch and pain (Esancy et al., 2018). While that study observed these effects using the weak TRPA1 activator imiquimod and the strong TRPA1 activator AITC, we see similar effects using varying concentrations of S1P (Fig. 1). It is interesting that our results in the cheek model mirror theirs, and lend support to a peripheral “population coding” model wherein low intensity stimuli can selectively drive pruriceptor activation and itch while high intensity stimuli activate both nociceptors and pruriceptors to drive pain and itch. We show that more S1P/S1PR3 signaling is required to activate nociceptors and drive pain behaviors than is required to activate pruriceptors and trigger itch.

But how, mechanistically, does this dose-dependent and selective activation of itch versus pain by S1P occur? A number of studies have generally implicated calcium-activated chloride and voltage-dependent potassium and sodium channels as being important for S1P-evoked neuronal excitation (Zhang et al., 2006a; Camprubi-Robles et al., 2013; Li et al., 2015). Here we show for the first time that both TRPA1 and TRPV1 play a key role in S1P-evoked AP firing in distinct subsets of somatosensory neurons and in itch and pain behaviors. TRP channels are nonselective cation channels that trigger calcium influx and depolarize the cell, and serve as important upstream transducers that trigger the activation of calcium- and voltage-sensitive channels. While S1P may activate multiple channels in somatosensory neurons, we have used pharmacological and genetic tools to show that TRPA1 and TRPV1 are essential for initiating S1P-evoked excitation and are differentially required for S1P-evoked itch and pain, respectively. Our experiments using G-protein pathway inhibitors suggest that S1PR3 utilizes distinct G-protein pathways to activate TRPA1 and TRPV1, which may contribute to differential initiation of itch and pain behaviors, and support a recent study suggesting that S1PR3 can couple to a number of downstream G-protein signaling pathways (Flock et al., 2017). While it is difficult to probe G-protein coupled pathways *in vivo,* it is tempting to speculate that the different signaling pathways downstream of S1PR3 *in vitro* may account for the differential engagement of pain and itch pathways by S1P.

While S1P signaling in general has been implicated in a variety of inflammatory skin diseases, our data suggest that S1PR3 may be an important contributor to cutaneous itch associated with these disorders. Fingolimod, which decreases S1PR activity at S1PRs 1,3,4 and 5 with varying affinity, has proven to be effective in reducing inflammation in mouse models of allergic contact dermatitis and in spontaneous dermatitis observed in the Nc/Nga mouse strain (Kohno et al., 2004; Sugita et al., 2010). Similarly, ponesimod, which inhibits S1PR1, decreases disease severity in psoriasis patients (D’ambrosio et al., 2016). In contrast, one study found that topical S1P can exert protective effects in mouse models of psoriasis and allergic contact dermatitis (Schaper et al., 2013). This is surprising, as one would expect S1P to promote inflammation, rather than inhibit it. However, this may be due to the diverse roles different S1PRs may play in different cell types. For example, S1PR2 exerts proliferative effects on keratinocytes (Japtok et al., 2014), whereas S1PR1 directs immune cell migration into tissues (Matloubian et al., 2004). Our findings suggest that S1P/S1PR3 signaling may play an important role in itch sensations associated with these skin disorders.

Several S1PR modulators are currently in use or in clinical trial for diseases linked to pain and/or itch, including multiple sclerosis, IBD, and psoriasis (Brinkmann et al., 2010; Kunkel et al., 2013; Degagné and Saba, 2014). Of particular note is ponesimod, which is in clinical trials for psoriasis treatment (Vaclavkova et al., 2014). Whereas ponesimod acts on the immune system and targets S1PR1 to combat inflammation, we propose neuronal S1P/S1PR3 signaling as a potential target for the treatment of inflammatory pain and itch. We showed previously that loss of S1PR3 can block inflammatory pain without affecting immune cell recruitment, and that a selective S1PR3 antagonist can ameliorate inflammatory hypersensitivity, suggesting that S1PR3-specific blockers may be effective for treating pain (Hill et al., 2018). Our finding that S1P can act as a pruritogen also suggests a role for S1P/S1PR3 in chronic itch, where circulating S1P levels have been found to be significantly increased in human psoriasis patients (Checa et al., 2015; Myśliwiec et al., 2016). While we and others have demonstrated the relevance of S1PR3 signaling to chronic pain, it will be essential to explore the role of S1P/S1PR3 signaling in chronic itch models to ascertain its pathological relevance. Our data supports a key role for S1PR3 in activation of pain and itch neurons and may inform rational design of S1P and S1PR modulators to treat pain, itch, and inflammatory skin diseases.

## Acknowledgements

We would like to thank Z. Rifi for assistance with scoring mouse behavior, N. Kucirek for critical reading of the manuscript, and all members of the D.M.B laboratory for constructive feedback and criticism.

## References

1. Akiyama T, Carstens E (2013) Neural processing of itch. Neuroscience 250:697–71 Available at: http://dx.doi.org/10.1016/j.neuroscience.2013.07.035.

2. Bandell M, Story GM, Hwang SW, Viswanath V, Eid SR, Petrus MJ, Earley TJ, Patapoutian A (2004) Noxious cold ion channel TRPA1 is activated by pungent compounds and bradykinin. Neuron 41:849–857.

3. Bäumer W, Rossbach K, Mischke R, Reines I, Langbein-Detsch I, Lüth A, Kleuser B (2011) Decreased concentration and enhanced metabolism of sphingosine-1-phosphate in lesional skin of dogs with atopic dermatitis: disturbed sphingosine-1-phosphate homeostasis in atopic dermatitis. J Invest Dermatol 131:266–268.

4. Bautista DM, Jordt SE, Nikai T, Tsuruda PR, Read AJ, Poblete J, Yamoah EN, Basbaum AI, Julius D (2006a) TRPA1 Mediates the Inflammatory Actions of Environmental Irritants and Proalgesic Agents. Cell 124:1269–1282.

5. Bautista DM, Jordt SE, Nikai T, Tsuruda PR, Read AJ, Poblete J, Yamoah EN, Basbaum AI, Julius D (2006b) TRPA1 Mediates the Inflammatory Actions of Environmental Irritants and Proalgesic Agents. Cell 124:1269–1282.

6. Brinkmann V, Billich A, Baumruker T, Heining P, Schmouder R, Francis G, Aradhye S, Burtin P (2010) Fingolimod (FTY720): discovery and development of an oral drug to treat multiple sclerosis. Nat Rev Drug Discov 9:883–897 Available at: http://www.ncbi.nlm.nih.gov/pubmed/21031003.

7. Brossard P, Derendorf H, Xu J, Maatouk H, Halabi A, Dingemanse J (2013) Pharmacokinetics and pharmacodynamics of ponesimod, a selective S1P 1 receptor modulator, in the first-in-human study. Br J Clin Pharmacol 76:888–896.

8. Camprubi-Robles M, Mair N, Andratsch M, Benetti C, Beroukas D, Rukwied R, Langeslag M, Proia RL, Schmelz M, Ferrer Montiel A V, Haberberger R V, Kress M, Camprubí-Robles M (2013) Sphingosine-1-phosphate-induced nociceptor excitation and ongoing pain behavior in mice and humans is largely mediated by S1P3 receptor. J Neurosci 33:2582–2592 Available at: http://www.ncbi.nlm.nih.gov/pubmed/23392686.

9. Castelino F V., Varga J (2014) Emerging cellular and molecular targets in fibrosis: Implications for scleroderma pathogenesis and targeted therapy. Curr Opin Rheumatol 26:607–614.

10. Caterina MJ, Leffler A, Malmberg AB, Martin WJ, Trafton J, Petersen-Zeitz KR, Koltzenburg M, Basbaum AI, Julius D (2000) Impaired Nociception and Pain Sensation in Mice Lacking the Capsaicin Receptor. Science (80-) 288:306–313 Available at: http://www.sciencemag.org/cgi/doi/10.1126/science.288.5464.306.

11. Checa A, Xu N, Sar DG, Haeggström JZ, Ståhle M, Wheelock CE (2015) Circulating levels of sphingosine-1-phosphate are elevated in severe, but not mild psoriasis and are unresponsive to anti-TNF-α treatment. Sci Rep 5:12017 Available at: http://www.ncbi.nlm.nih.gov/pubmed/26174087.

12. Chiba Y, Takeuchi H, Sakai H, Misawa M (2010) SKI-II, an inhibitor of sphingosine kinase, ameliorates antigen-induced bronchial smooth muscle hyperresponsiveness, but not airway inflammation, in mice. J Pharmacol Sci 114:304–310.

13. Choi JW, Gardell SE, Herr DR, Rivera R, Lee C-W, Noguchi K, Teo ST, Yung YC, Lu M, Kennedy G, Chun J (2011) FTY720 (fingolimod) efficacy in an animal model of multiple sclerosis requires astrocyte sphingosine 1-phosphate receptor 1 (S1P1) modulation. Proc Natl Acad Sci U S A 108:751–756 Available at: papers3://publication/doi/10.1073/pnas.1014154108/-/DCSupplemental/pnas.201014154SI.pdf%5Cnhttp://www.pubmedcentral.nih.gov/articlerender.fcgi?artid=3021041&tool=pmcentrez&rendertype=abstract.

14. Chuang HH, Prescott ED, Kong H, Shields S, Jordt SE, Basbaum a I, Chao M V, Julius D (2001) Bradykinin and nerve growth factor release the capsaicin receptor from PtdIns(4,5)P2-mediated inhibition. Nature 411:957–962.

15. Cyster JG, Schwab SR (2012) Sphingosine-1-Phosphate and Lymphocyte Egress from Lymphoid Organs. Annu Rev Immunol 30:69–94 Available at: http://www.annualreviews.org/doi/abs/10.1146/annurev-immunol-020711-075011.

16. D’ambrosio D, Freedman MS, Prinz J (2016) Ponesimod, a selective S1P1 receptor modulator: A potential treatment for multiple sclerosis and other immune-mediated diseases. Ther Adv Chronic Dis 7:18–33.

17. Dai Y, Wang S, Tominaga M, Yamamoto S, Fukuoka T, Higashi T, Kobayashi K, Obata K, Yamanaka H, Noguchi K (2007) Sensitization of TRPA1 by PAR2 contributes to the sensation of inflammatory pain. J Clin Invest 117:1979–1987.

18. Degagné E, Saba JD (2014) S1 pping fire: Sphingosine-1-phosphate signaling as an emerging target in inflammatory bowel disease and colitis-associated cancer. Clin Exp Gastroenterol 7:205–214.

19. Donoviel MS, Hait NC, Ramachandran S, Maceyka M, Takabe K, Milstien S, Oravecz T, Spiegel S (2015) Spinster 2, a sphingosine 1-phosphate transporter, plays a Critical Role in inflammatory and Autoimmune Diseases. FASEB J 29:5018–5028.

20. Esancy K, Condon L, Feng J, Kimball C, Curtright A, Dhaka A (2018) A zebrafish and mouse model for selective pruritus via direct activation of TRPA1. Elife 7:1–24 Available at: http://www.ncbi.nlm.nih.gov/pubmed/29561265%0Ahttps://elifesciences.org/articles/32036.

21. Finley a, Chen Z, Esposito E, Cuzzocrea S, Sabbadini R, Salvemini D (2013) Sphingosine 1-phosphate mediates hyperalgesia via a neutrophil-dependent mechanism. PLoS One 8:e55255 Available at: http://www.ncbi.nlm.nih.gov/pubmed/23372844%5Cnhttp://www.plosone.org/article/fetchObject.action?uri=info:doi/10.1371/journal.pone.0055255&representation=PDF.

22. Flock T, Hauser AS, Lund N, Gloriam DE, Balaji S, Babu MM (2017) Selectivity determinants of GPCR-G-protein binding. Nature 545:317–322 Available at: http://dx.doi.org/10.1038/nature22070.

23. Gao Y, Cao E, Julius D, Cheng Y (2016) TRPV1 structures in nanodiscs reveal mechanisms of ligand and lipid action. Nature 534:347–351 Available at: http://dx.doi.org/10.1038/nature17964.

24. Gerhold K a., Bautista DM (2009) Molecular and cellular mechanisms of trigeminal chemosensation. Ann N Y Acad Sci 1170:184–189.

25. Gerhold K a., Pellegrino M, Tsunozaki M, Morita T, Leitch DB, Tsuruda PR, Brem RB, Catania KC, Bautista DM (2013) The Star-Nosed Mole Reveals Clues to the Molecular Basis of Mammalian Touch. PLoS One 8.

26. Grenald SA, Doyle TM, Zhang H, Slosky LM, Chen Z, Largent-Milnes TM, Spiegel S, Vanderah TW, Salvemini D (2017) Targeting the S1P/S1PR1 axis mitigates cancer-induced bone pain and neuroinflammation. Pain 158:1 Available at: http://insights.ovid.com/crossref?an=00006396-900000000-99231%5Cnhttp://www.ncbi.nlm.nih.gov/pubmed/28570482.

27. Han L, Ma C, Liu Q, Weng H, Cui Y, Tang Z, Guan Y, Xiao B, Lamotte R, Dong X (2013) A subpopulation of nociceptors specifically linked to itch. Nat Neurosci 16:174–182.

28. Hill RZ, Hoffman BU, Morita T, Campos SM, Lumpkin EA, Brem RB, Bautista DM (2018) The signaling lipid sphingosine 1-phosphate regulates mechanical pain. Elife 7:e33285 Available at: https://elifesciences.org/articles/33285.

29. Hou J-C, Fang X-M (2015) S1P-S1PRs Alliance in Neuropathic Pain Processing. J Anaesth Perioper Med 2:1–9.

30. Idzko M, Panther E, Corinti S, Morelli A, Ferrari D, Herouy Y, Dichmann S, Mockenhaupt M, Gebicke-Haerter P, Di Virgilio F, Girolomoni G, Norgauer J (2002) Sphingosine 1-phosphate induces chemotaxis of immature and modulates cytokine-release in mature human dendritic cells for emergence of Th2 immune responses. FASEB J 16:625–627.

31. Imamachi N, Park GH, Lee H, Anderson DJ, Simon MI, Basbaum AI, Han S-K (2009) TRPV1-expressing primary afferents generate behavioral responses to pruritogens via multiple mechanisms. Proc Natl Acad Sci U S A 106:11330–11335.

32. Janes K, Little JW, Li C, Bryant L, Chen C, Chen Z, Kamocki K, Doyle T, Snider A, Esposito E, Cuzzocrea S, Bieberich E, Obeid L, Petrache I, Nicol G, Neumann WL, Salvemini D (2014) The development and maintenance of paclitaxel-induced neuropathic pain require activation of the sphingosine 1-phosphate receptor subtype 1. J Biol Chem 289:21082–21097.

33. Japtok L, Bäumer W, Kleuser B (2014) Sphingosine-1-phosphate as signaling molecule in the skin. Allergo J Int 23:54–59 Available at: http://link.springer.com/10.1007/s40629-014-0008-2.

34. Kim E-S, Kim J-S, Kim SG, Hwang S, Lee CH, Moon A (2011) Sphingosine 1-phosphate regulates matrix metalloproteinase-9 expression and breast cell invasion through S1P3-G q coupling. J Cell Sci 124:2220–2230 Available at: http://jcs.biologists.org/cgi/doi/10.1242/jcs.076794.

35. Kohno T, Tsuji T, Hirayama K, Watabe K, Matsumoto A, Kohno T, Fujita T (2004) A novel immunomodulator, FTY720, prevents spontaneous dermatitis in NC/Nga mice. Biol Pharm Bull 27:1392–1396 Available at: http://www.ncbi.nlm.nih.gov/pubmed/15340225.

36. Kono M, Mi Y, Liu Y, Sasaki T, Allende ML, Wu YP, Yamashita T, Proia RL (2004) The sphingosine-1-phosphate receptors S1P1, S1P2, and S1P3 function coordinately during embryonic angiogenesis. J Biol Chem 279:29367–29373.

37. Krause A, D’Ambrosio D, Dingemanse J (2017) Modeling clinical efficacy of the S1P receptor modulator ponesimod in psoriasis. J Dermatol Sci 89:136–145 Available at: http://dx.doi.org/10.1016/j.jdermsci.2017.11.003.

38. Kunkel GT, Maceyka M, Milstien S, Spiegel S (2013) Targeting the sphingosine-1-phosphate axis in cancer, inflammation and beyond. Nat Rev Drug Discov 12:688–702 Available at: http://www.pubmedcentral.nih.gov/articlerender.fcgi?artid=3908769&tool=pmcentrez&rendertype=abstract.

39. Kwon Y, Shim H-S, Wang X, Montell C (2008) Control of thermotactic behavior via coupling of a TRP channel to a phospholipase C signaling cascade. Nat Neurosci 11:871–873.

40. Li C, Li J, Kays J, Guerrero M, Nicol GD (2015) Sphingosine 1-phosphate enhances the excitability of rat sensory neurons through activation of sphingosine 1-phosphate receptors 1 and/or 3. J Neuroinflammation 12:1–20 Available at: http://www.jneuroinflammation.com/content/12/1/70.

41. Liang J, Nagahashi M, Kim EY, Harikumar KB, Yamada A, Huang WC, Hait NC, Allegood JC, Price MM, Avni D, Takabe K, Kordula T, Milstien S, Spiegel S (2013) Sphingosine-1-Phosphate Links Persistent STAT3 Activation, Chronic Intestinal Inflammation, and Development of Colitis-Associated Cancer. Cancer Cell 23:107–120 Available at: http://dx.doi.org/10.1016/j.ccr.2012.11.013.

42. Liu Q, Sikand P, Ma C, Tang Z, Han L, Li Z, Sun S, LaMotte RH, Dong X (2012) Mechanisms of itch evoked by β-alanine. J Neurosci 32:14532–14537 Available at: http://www.pubmedcentral.nih.gov/articlerender.fcgi?artid=3491570%7B&%7Dtool=pmcentrez%7B&%7Drendertype=abstract.

43. Liu Q, Tang Z, Surdenikova L, Kim S, Patel KN, Kim A, Ru F, Guan Y, Weng HJ, Geng Y, Undem BJ, Kollarik M, Chen ZF, Anderson DJ, Dong X (2009) Sensory Neuron-Specific GPCR Mrgprs Are Itch Receptors Mediating Chloroquine-Induced Pruritus. Cell 139:1353–1365 Available at: http://dx.doi.org/10.1016/j.cell.2009.11.034.

44. Mair N, Benetti C, Andratsch M, Leitner MG, Constantin CE, Camprubí-Robles M, Quarta S, Biasio W, Kuner R, Gibbins IL, Kress M, Haberberger R V. (2011) Genetic evidence for involvement of neuronally expressed s1P1 receptor in nociceptor sensitization and inflammatory pain. PLoS One 6.

45. Matloubian M, Lo CG, Cinamon G, Lesneski MJ, Xu Y, Brinkmann V, Allende ML, Proia RL, Cyster JG (2004) Lymphocyte egress from thymus and peripheral lymphoid organs is dependent on S1P receptor 1. Nature 427:355–360.

46. Moore C, Gupta R, Jordt S-E, Chen Y, Liedtke WB (2017) Regulation of Pain and Itch by TRP Channels. Neurosci Bull Available at: http://www.ncbi.nlm.nih.gov/pubmed/29282613%0Ahttp://link.springer.com/10.1007/s12264-017-0200-8.

47. Morita T, McClain SP, Batia LM, Pellegrino M, Wilson SR, Kienzler MA, Lyman K, Olsen ASB, Wong JF, Stucky CL, Brem RB, Bautista DM (2015) HTR7 Mediates Serotonergic Acute and Chronic Itch. Neuron 87:124–138 Available at: http://dx.doi.org/10.1016/j.neuron.2015.05.044.

48. Myśliwiec H, Baran A, Harasim-Symbor E, Choromańska B, Myśliwiec P, Milewska AJ, Chabowski A, Flisiak I (2016) Increase in circulating sphingosine-1-phosphate and decrease in ceramide levels in psoriatic patients. Arch Dermatol Res Available at: http://www.ncbi.nlm.nih.gov/pubmed/27988894%0Ahttp://link.springer.com/10.1007/s00403-016-1709-9.

49. Olivera A, Allende ML, Proia RL (2013) Shaping the landscape: Metabolic regulation of S1P gradients. Biochim Biophys Acta - Mol Cell Biol Lipids 1831:193–202 Available at: http://dx.doi.org/10.1016/j.bbalip.2012.06.007.

50. Patti G, Yanes O, Shriver L, Courade J-P, Tautenhahn R, Manchester M, Siuzdak G (2008) Metabolomics implicates altered sphingolipids in chronic pain of neuropathic origin. Nat Chem Biol 8:232–234.

51. Paulsen CE, Armache J, Gao Y, Cheng Y, Julius D (2015) Structure of the TRPA1 ion channel suggests regulatory mechanisms. Nature 520:511–517.

52. Prescott E, Julius D (2003) A Modular PIP 2 Binding Site as a Determinant of Capsaicin Receptor Sensitivity. Science (80-) 300:1284–1288.

53. Reddy VB, Sun S, Azimi E, Elmariah SB, Dong X, Lerner EA (2015) Redefining the concept of protease-activated receptors: cathepsin S evokes itch via activation of Mrgprs. Nat Commun 6:7864 Available at: http://www.nature.com/ncomms/2015/150728/ncomms8864/full/ncomms8864.html.

54. Rivera J, Proia RL, Olivera A (2008) The alliance of sphingosine-1-phosphate and its receptors in immunity. Nat Rev Immunol 8:753–763 Available at: http://www.nature.com/doifinder/10.1038/nri2400.

55. Rogerio AP, Andrade EL, Calixto JB (2011) C-fibers, but not the transient potential receptor vanilloid 1 (TRPV1), play a role in experimental allergic airway inflammation. Eur J Pharmacol 662:55–62 Available at: http://dx.doi.org/10.1016/j.ejphar.2011.04.027.

56. Rohacs T, Thyagarajan B, Lukacs V (2008) Phospholipase C mediated modulation of TRPV1 channels. Mol Neurobiol 37:153–163.

57. Roviezzo F, Sorrentino R, Bertolino A, De Gruttola L, Terlizzi M, Pinto A, Napolitano M, Castello G, D’Agostino B, Ianaro A, Sorrentino R, Cirino G (2015) S1P-induced airway smooth muscle hyperresponsiveness and lung inflammation in vivo: Molecular and cellular mechanisms. Br J Pharmacol 172:1882–1893.

58. Saluja R, Kumar A, Jain M, Goel SK, Jain A (2017) Role of sphingosine-1-phosphate in mast cell functions and asthma and its regulation by non-coding RNA. Front Immunol 8:1–7.

59. Sanna MG, Vincent KP, Repetto E, Nhan N, Brown SJ, Abgaryan L, Riley SW, Leaf NB, Cahalan SM, Kiosses WB, Kohno Y, Heller Brown J, McCulloch AD, Rosen H, Gonzalez-Cabrera PJ (2016) Bitopic S1P3 Antagonist Rescue from Complete Heart Block: Pharmacological and Genetic Evidence for Direct S1P3 Regulation of Mouse Cardiac Conduction. Mol Pharmacol 89:176–186 Available at: http://molpharm.aspetjournals.org/cgi/doi/10.1124/mol.115.100222.

60. Schaper K, Dickhaut J, Japtok L, Kietzmann M, Mischke R, Kleuser B, Bäumer W (2013) Sphingosine-1-phosphate exhibits anti-proliferative and anti-inflammatory effects in mouse models of psoriasis. J Dermatol Sci 71:29–36.

61. Schmidt M, Dubin AE, Petrus MJ, Earley TJ, Patapoutian A (2009) Nociceptive Signals Induce Trafficking of TRPA1 to the Plasma Membrane. Neuron 64:498–509 Available at: http://dx.doi.org/10.1016/j.neuron.2009.09.030.

62. Schüppel M, Kürschner U, Kleuser U, Schäfer-Korting M, Kleuser B (2008) Sphingosine 1-phosphate restrains insulin-mediated keratinocyte proliferation via inhibition of Akt through the S1P2 receptor subtype. J Invest Dermatol 128:1747–1756.

63. Shim W-S, Tak M-H, Lee M-H, Kim M, Kim M, Koo J-Y, Lee C-H, Kim M, Oh U (2007) TRPV1 mediates histamine-induced itching via the activation of phospholipase A2 and 12-lipoxygenase. J Neurosci 27:2331–2337 Available at: http://www.jneurosci.org/cgi/content/full/27/9/2331.

64. Shimada SG, LaMotte RH (2008) Behavioral differentiation between itch and pain in mouse. Pain 139:681–687.

65. Sugita K, Kabashima K, Sakabe JI, Yoshiki R, Tanizaki H, Tokura Y (2010) FTY720 regulates bone marrow egress of eosinophils and modulates late-phase skin reaction in mice. Am J Pathol 177:1881–1887 Available at: http://dx.doi.org/10.2353/ajpath.2010.100119.

66. Thieme M, Zillikens D, Sadik CD (2017) Sphingosine-1-phosphate modulators in inflammatory skin diseases – lining up for clinical translation. Exp Dermatol 26:206–210.

67. Trankner D, Hahne N, Sugino K, Hoon MA, Zuker C (2014) Population of sensory neurons essential for asthmatic hyperreactivity of inflamed airways. Proc Natl Acad Sci 111:11515–11520 Available at: http://www.pnas.org/cgi/doi/10.1073/pnas.1411032111.

68. Tsunozaki M, Lennertz RC, Vilceanu D, Katta S, Stucky CL, Bautista DM (2013) A “toothache tree” alkylamide inhibits Aδ mechanonociceptors to alleviate mechanical pain. J Physiol 591:3325–3340 Available at: http://jp.physoc.org/content/591/13/3325.long.

69. Usoskin D, Furlan A, Islam S, Abdo H, Lönnerberg P, Lou D, Hjerling-Leffler J, Haeggström J, Kharchenko O, Kharchenko P V, Linnarsson S, Ernfors P (2015) Unbiased classification of sensory neuron types by large-scale single-cell RNA sequencing. Nat Neurosci 18:145–153 Available at: http://dx.doi.org/10.1038/nn.3881.

70. Vaclavkova A, Chimenti S, Arenberger P, Holló P, Sator PG, Burcklen M, Stefani M, D’Ambrosio D (2014) Oral ponesimod in patients with chronic plaque psoriasis: A randomised, double-blind, placebo-controlled phase 2 trial. Lancet 384:2036–2045.

71. Wang J, Kollarik M, Ru F, Sun H, Mcneil B, Dong X, Stephens G, Korolevich S, Brohawn P, Kolbeck R, Undem B (2017) Distinct and common expression of receptors for inflammatory mediators in vagal nodose versus jugular capsaicin-sensitive / TRPV1-positive neurons detected by low input RNA sequencing. :1–20.

72. Weth D, Benetti C, Rauch C, Gstraunthaler G, Schmidt H, Geisslinger G, Sabbadini R, Proia RL, Kress M (2015) Activated platelets release sphingosine 1-phosphate and induce hypersensitivity to noxious heat stimuli in vivo. Front Neurosci 9:1–8 Available at: http://journal.frontiersin.org/article/10.3389/fnins.2015.00140/abstract.

73. Wilson SR, Gerhold K a, Bifolck-Fisher A, Liu Q, Patel KN, Dong X, Bautista DM (2011) TRPA1 is required for histamine-independent, Mas-related G protein-coupled receptor-mediated itch. Nat Neurosci 14:595–602 Available at: http://dx.doi.org/10.1038/nn.2789.

74. Yamashita-Sugahara Y, Tokuzawa Y, Nakachi Y, Kanesaki-Yatsuka Y, Matsumoto M, Mizuno Y, Okazaki Y (2013) Fam57b (Family with sequence similarity 57, member B), a novel peroxisome proliferator-activated receptor y target gene that regulates adipogenesis through ceramide synthesis. J Biol Chem 288:4522–4537.

75. Zhang YH, Fehrenbacher JC, Vasko MR, Nicol GD (2006a) Sphingosine-1-Phosphate Via Activation of a G-Protein-Coupled Receptor(s) Enhances the Excitability of Rat Sensory Neurons. J Neurophysiol 96:1042–1052 Available at: http://www.physiology.org/doi/10.1152/jn.00120.2006.

76. Zhang YH, Vasko MR, Nicol GD (2006b) Intracellular sphingosine 1-phosphate mediates the increased excitability produced by nerve growth factor in rat sensory neurons. J Physiol 575:101–113.

77. Zhu Y, Hanson CE, Liu Q, Han L (2017) Mrgprs activation is required for chronic itch conditions in mice. Itch 2:e09.

